# A thermodynamic chromatin polymer model characterizes the epigenetic conditions for *Hox* collinearity

**DOI:** 10.64898/2025.12.28.696739

**Authors:** Yoshifumi Asakura, Yoshihiro Morishita, Kyosuke Adachi, Takayuki Suzuki

## Abstract

*Hox* genes consist of four clusters, A, B, C, and D, each containing up to 13 paralogous genes. They are linearly arrayed on cluster regions in the same order as their expression timings along the anterior-posterior (A-P) axis development. This correspondence is known as temporal collinearity, and when the expression order of *Hox* genes is disrupted, the A-P axis is not accurately established. Although essential molecules and genomic regulatory elements for *Hox* collinearity have been identified, it remains unclear whether they act on individual genes or instead coordinate the cluster-wide sequential activation along the genomic order. To address this question, we developed a theoretical framework based on the statistical thermodynamics of chromatin modeled as a polymer, interpreting collinearity as a sequential opening of nucleosomes along the gene cluster. The model incorporates interactions between nucleosomes and DNA-binding factors, including transcription factors, represented by energy terms for binding and chromatin bridging. This formulation links the thermodynamic stability of chromatin states with the sequential opening process along the *Hox* clusters. Our model characterized key quantities that determine the order of chromatin opening and revealed intrinsic constraints that limit sequential opening of chromatin, underlying *Hox* collinearity.

**Significance statement:** *Hox* collinearity, a remarkable correspondence between the genomic order and expression timing of *Hox* genes, provides cells with positional information on the anterior-posterior axis during development. Although essential molecules and genomic regulatory elements have been identified, the mechanism of sequential activation along the genomic order remains elusive. To address this question, we theoretically investigated the epigenetic states of chromatin using a chromatin polymer model in statistical thermodynamics. Our model provides a possible explanation for the sequential activation of the chromatin underlying *Hox* collinearity. This work bridges developmental genetics and statistical thermodynamics, offering a quantitative framework for understanding gene activation order along the genome.

## 1 Introduction

Epigenetics is an essential element in the regulation of gene expression. It was originally defined as stable and heritable changes in gene expression patterns without any DNA sequence alterations [1, 2], and is known to involve DNA methylation, histone modifications, and three-dimensional (3D) chromatin organization [3–12]. Since the development of the Hi-C technique [13], the regulatory significance of 3D chromatin organization has been quantitatively demonstrated throughout the genome in many species [12, 14, 15].

Transcription factors (TFs) are DNA-binding factors that regulate gene expression. TFs bind to cis-regulatory elements (CREs), such as promoters and enhancers. Enhancers can be located far from their target genes, especially in vertebrates. Such binding events mediate enhancer-promoter proximity, which is essential for the transcriptional activation of genes [16–19]. TFs also induce biochemical reactions in chromatin, such as demethylation, by recruiting ten-eleven translocation (TET) proteins [15, 20–23]. Because of this molecular system of transcription, 3D geometries, chromatin state, and their interplay are important for understanding gene expression regulation.

Among many transcriptional regulation systems, that of the *Hox* genes is one of the most intriguing and puzzling systems. The *Hox* genes consist of four clusters, A, B, C, and D, each comprising a series of up to 13 paralogous genes [24–27]. They are linearly arrayed within each cluster in the same order as their expression timings during anterior-posterior (A-P) axis development [27–29]. This correspondence is known as temporal collinearity [29]. Because vertebrate development progresses from anterior to posterior, the order of the *Hox* genes on genome also corresponds to the spatial domains of expression [30]. This correspondence is known as spatial collinearity [29, 31]. *Hox* genes encode TFs [32], and their expression patterns provide cells with positional information on the A-P axis during development [33]. When the temporal or spatial order of *Hox* gene expression is disrupted, the A-P axis is not accurately established [34–37]. Enhancers located near the *Hox* clusters have been reported to regulate their expression [38–41]. Additionally, several secreted signaling molecules — including growth and differentiation factor-11 (GDF11), a member of the TGF-*β* superfamily, WNT ligands, and retinoic acid — have been shown to regulate *Hox* gene expression [37, 42, 43].

Although essential molecules and CREs for *Hox* collinearity have been identified, it remains unclear whether they act on individual genes or instead coordinate the cluster-wide sequential activation along the genomic order [27]. Genomic position effects on expression have also been reported beyond the *Hox* loci [44]. For example, some enhancers simultaneously activate multiple neighboring genes [45, 46]. Additionally, it has been reported that a highly active locus can stimulate transcription at neighboring loci [47]. *β*-globin genes are also expressed in the same order on the *β*-globin gene cluster during development [48, 49]. Despite the regulatory importance of position effects [50, 51] and enhancer sharing [45, 46], the mechanisms by which clustered genes achieve collinearity have remained an open question [27].

To address the mechanisms of cluster-wide sequential activation of genes, here we theoretically investigated chromatin conformations and dynamics, building on recent studies in statistical thermodynamics which have shown that chromatin conformations and dynamics can be recapitulated by polymer models [11, 52–54]. We interpreted collinearity as a phenomenon in which nucleosomes open sequentially along the cluster. To investigate conditions for sequential opening, we first defined a chromatin model as a polymer that interacts with TFs and other DNA-binding factors. Next, we analyzed the energy change during the opening of a nucleosome on the chromatin and identified a relationship between the chromatin opening order and epigenetic conditions. This analysis revealed key quantities that determine the chromatin opening order and intrinsic constraints in achieving sequential chromatin opening along the gene order on the genome. Our approach, analyzing the directionality of epigenetic state changes by a chromatin polymer model, established a framework that identifies the relationship between epigenetics and its dynamics.

## 2 Results

### 2.1 A polymer model with DNA-binding factor interaction

Here, we interpreted collinearity as a phenomenon in which nucleosomes open sequentially. To investigate the nucleosome opening order along the genome, we modeled the chromatin as a linear polymer. In this model, the chromatin polymer consists of nucleosomes as monomers. Nucleosomes interact with each other, and TFs and other DNA-binding factors mediate these interactions (figure 1 a-d). These interactions were formulated in terms of an energy function. We analyzed the directionality of the nucleosome-state dynamics by evaluating the energy differences associated with state transitions. To formulate the energy, we defined a set of positions {**R**_*n*_} that represents the chromatin 3D coordination and a set of nucleosome states {*s*_*n*_} that represents the chemical modification of histones in each nucleosome. Here, we refer to the combination of {**R**_*n*_} and {*s*_*n*_} as the model epigenetic state. Histones generally undergo chemical modifications. Such modifications regulate the accessibility of various DNA-binding factors to DNA. For example, H3K4me3 and H3K27ac are enriched in active, accessible, and open regions. H3K27me3 and H3k9me3 are enriched in inactive, silent, and closed regions. To focus on the conditions of sequential changes of epigenetic states, especially from inactive-and-closed to the active-and-open states, we simplified the nucleosome states to open or closed hereafter. This simplification is supported by previous studies showing that chromatin dynamics can be sufficiently captured by two-state analysis [53, 54]. In the model, we denote *s*_*n*_ = 1 for the open state, and *s*_*n*_ = −1 for the closed state (a generalized formulation allowing multiple states is given in supplementary information), and the effects of the monomer states on the dynamics of the epigenetic state are parameterized by interaction coefficients as described later. Starting from a homopolymer model, which does not consider the states of the monomers, we added the effects of each monomer nucleosome state as,

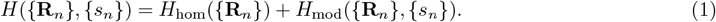

**Figure 1.**
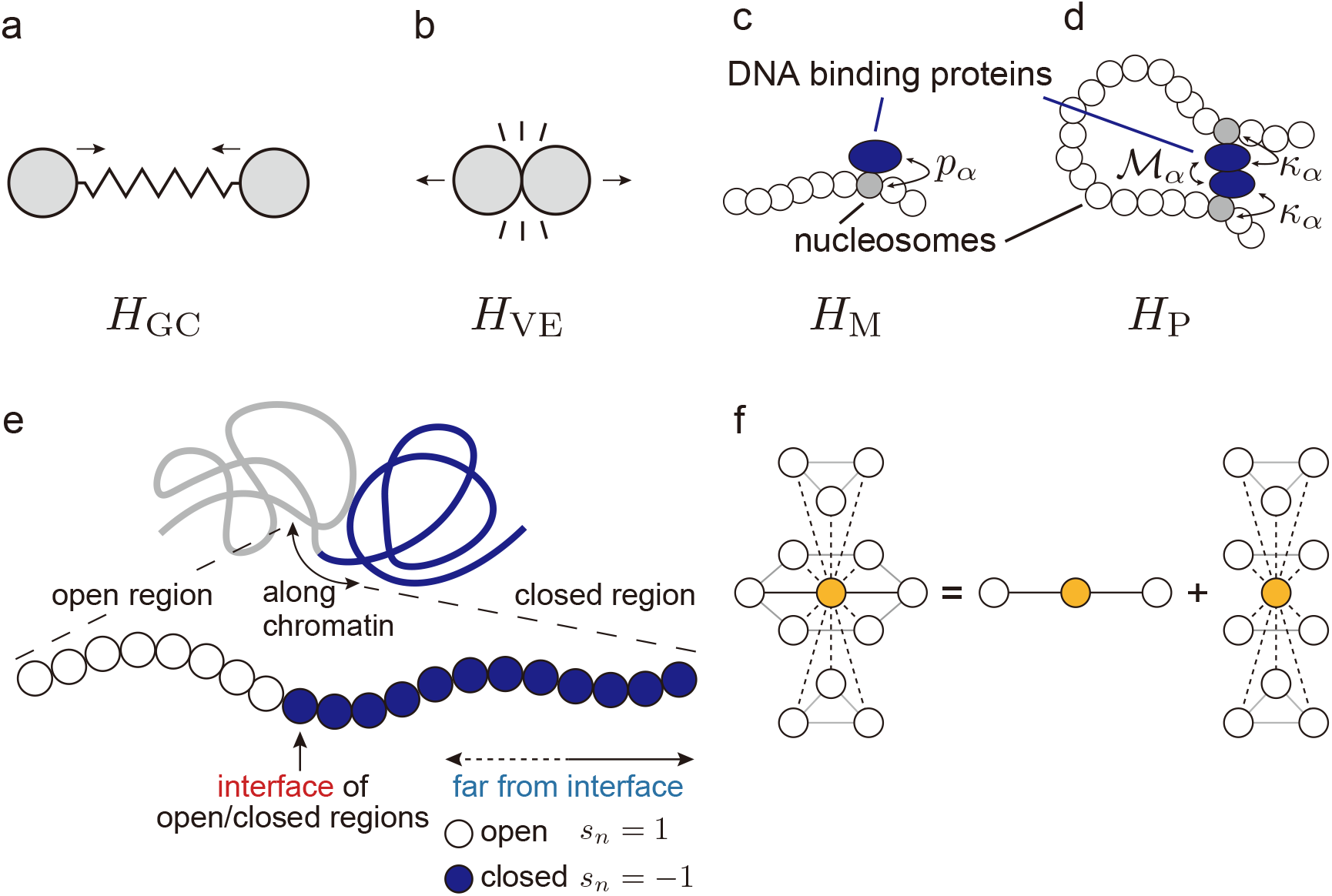
Model settings. (a-d) A model of polymer dynamics. It consists of (a) Gaussian Chain, (b) Volume Exclusion, (c) Mark-dependency, where *p*_*α*_ denotes the binding parameter, and (d) Pair effects, where *κ*_*α*_ and *M*_*α*_ denote the binding parameters between nucleosome and DNA-binding proteins. (e) An epigenetic state during sequential opening as an initial condition. Conditions in which the sequential opening continues or breaks down were analyzed. (f) Hexagonal closest packing. Particles with a common diameter can contact 12 particles at the maximum density. In the polymer case, there are two bonded neighbors (one-dimensional neighbors) and ten contacts with monomers by three-dimensional access.

The homopolymer model Hamiltonian reads

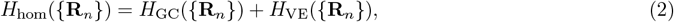

where *H*_GC_ is the polymer chain elasticity (i.e., the Gaussian chain model) (figure 1 a), and *H*_VE_ represents the volume exclusion effects (figure 1 b). These are independent of the nucleosome states. To capture the epigenetic regulations of the chromatin, we added histone modification-dependent energy *H*_mod_({**R**_*n*_}, {*s*_*n*_}) to our model. The Hamiltonian reads,

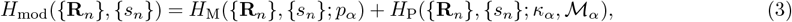

where *p*_*α*_, *κ*_*α*_, and ℳ_*α*_ are parameters as defined below (see supplementary information for details on mathematical settings). In the *H*_mod_({**R**_*n*_}, {*s*_*n*_}), *H*_M_ is the energy that one DNA-binding factor binds to one nucleosome, parameterized by a DNA-binding affinity constant *p*_*α*_ between a DNA-binding factor and a nucleosome (figure 1 c). *H*_P_ is the energy of pairing effects, in which two nucleosomes form a contact mediated by DNA-binding factors (figure 1 d). *H*_P_ is parameterized by two binding affinity constants, *κ*_*α*_ between a nucleosome and a DNA-binding factor *α*, and ℳ_*αβ*_ between two DNA-binding factors *α* and *β* binding to different nucleosomes respectively. When a pair of DNA-binding factors has binding parameters *κ*_*α*_*κ*_*β*_ ℳ_*αβ*_ > 0, it stabilizes interactions between nucleosomes in the same state. Heterochromatin protein 1 (HP1) [55, 56], Polycomb repressive complex 2 (PRC2) [57], and many TFs [16–19, 58] fit into this. In contrast, when a pair of DNA-binding factors has *κ*_*α*_*κ*_*β*_ ℳ_*αβ*_ < 0, it stabilizes interactions between nucleosomes in different states. However, as long as we surveyed, no such factors have been identified. Moreover, because many TFs have binding motifs both in enhancers and promoters of their target genes [59–64], one type of TF can mediate the interactions between CREs. Based on these biological tendency, we hereafter focus on

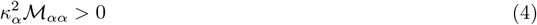

case. For *α*≠ *β* or *κ*_*α*_*κ*_*β*_ℳ _*αβ*_ < 0 case, see supplementary information. It is worth noting that the difference between *H*_M_ and *H*_P_ corresponds to the differences in DNA-binding energies between TFs and other DNA-binding factors. TFs form a loop-shaped bridge between two regions, and *H*_P_ models the bridging effect on the chromatin polymer. However, other DNA-binding factors do not necessarily form loop-shaped bridges. Reflecting this difference, all DNA-binding factors give *H*_M_ as nonzero values, and only TFs give *H*_P_ as nonzero values. Additionally, here we assumed that the monomers are spheres with nonzero radii instead of points without volumes. Assuming the closest packing in 3D space, one nucleosome can interact with up to 12 nucleosomes, including its bonded monomers (figure 1 f).

Next we derived energy inequalities that characterize the condition where sequential opening is favored, under the initial condition (figure 1 e) where an open region (i.e., an array of open nucleosomes) is connected to a closed region (i.e., an array of closed nucleosomes). At the boundary between the two regions, the interface nucleosome lies between an open nucleosome and a closed nucleosome along the 1D polymer. Other nucleosomes have two nucleosomes in the same state as their 1D neighbors on the polymer. In addition to the 1D interactions, there are 3D interactions, as described in *H*_VE_ and *H*_P_. In the sequential opening process, the interface nucleosome opens, while any nucleosomes in the closed region remain closed. This yields locally the same condition as the initial condition (figure 1 e) by regarding the next boundary nucleosome as the interface. Once any nucleosome in the closed region opens, it yields a condition different from the initial condition (figure 1 e). This represents a breakdown of the sequential opening process. Here we refer to this breakdown of the sequential opening process as random opening. To distinguish the sequential and random openings, we defined 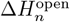, the energy difference between the open and closed states of the *n*th nucleosome,

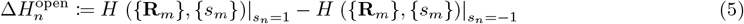

Since *H*_hom_({**R**_*n*_}) is independent of states {*s*_*n*_}, the difference in energy was derived as,

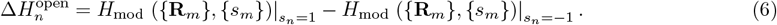

The sequential opening is formulated as the following two conditions:

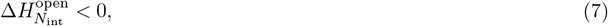

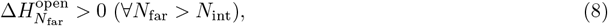

where *N*_int_ and *N*_far_ are indices of the interface nucleosome and a nucleosome far from the interface, respectively. If the inequality (7) does not hold, the interface nucleosome remains closed. If the inequality (8) does not hold, the sequential opening breaks down. Thus, the sequential opening condition was formulated by the inequalities (7) and (8).

### 2.2 Opening determinant: a criterion of nucleosome state transition

Next, we analyzed the energy difference 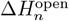, which governs the sequential opening condition formulated above. From the Hamiltonian (equations (1)-(3); derivation in supplementary information), we obtained an expression of the energy change:

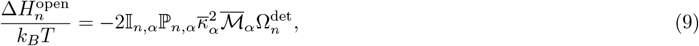

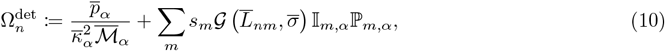

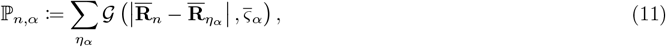

Where 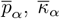, and 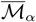 are dimensionless counterparts of the binding constants introduced in *H*_M_ and *H*_P_, scaled by *k*_*B*_*T* and reference volume (detailed derivation in supplementary information); 𝕀_*n,α*_ is a fraction of the target sites of the DNA-binding factor *α* within the polymer; ℙ_*n,α*_ is the mean concentration of a DNA-binding factor *α*; 𝒢 is a Gaussian distribution function; and 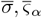 are dimensionless radii of a nucleosome and a DNA-binding factor *α*, respectively. In equation (9), the coefficients 𝕀_*n,α*_ and ℙ_*n,α*_ are positive constants, and here we assume 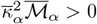 as discussed above. Therefore, we defined an opening determinant as equation (10), which determines the sign of energy change at the *n*th nucleosome. Thus, sequential opening (inequalities (7) and (8)) reduces to

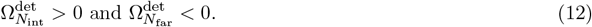

To investigate the key quantities that determine 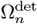, we applied an approximation by separating 1D and 3D contacts and replacing interaction terms with their averaged effects. Because a nucleosome is a monomer on a chromatin polymer without branches, there are only two nucleosomes indexed by *n* − 1 and *n* + 1 beside *n* on a 1D chromatin polymer. Other nucleosomes can interact with the *n*th nucleosome by 3D contacts. Therefore, 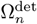 was divided into 1D and 3D effects as,

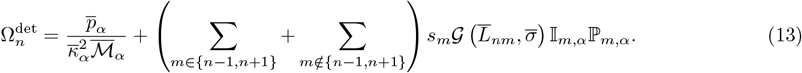

By averaging the interaction effects of contacts around the *n*th nucleosome, separately for 1D and 3D interactions, 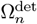 was approximated (see the supplementary information for the details) as a function of four parameters:

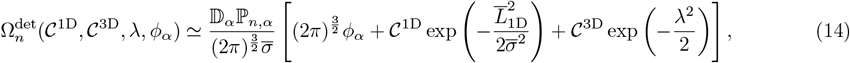

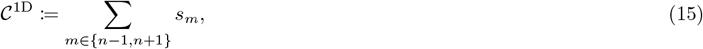

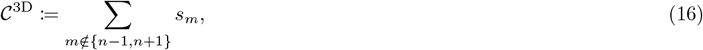

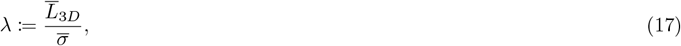

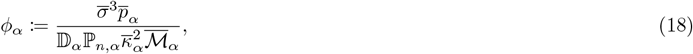

Where 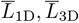, and 𝔻_*α*_ are the dimensionless distances from the *n*th nucleosome to its 1D neighbors, to its 3D neighbors, and an approximated fraction of target sites 𝕀_*n,α*_ of the DNA-binding factor *α*, respectively. 𝒞^1D^ and 𝒞^3D^ are the effects of contact with 1D and 3D neighbors, respectively. Through the parameters 𝒞^1D^ and 𝒞^3D^, states {*s*_*n*_} affect 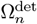. 𝒞^1D^ reflects the local states around the *n*th nucleosome. When both 1D neighbors of the *n*th nucleosome are open, 𝒞^1D^ = 2. When both are closed, 𝒞^1D^ = −2. When the *n*th nucleosome is between an open and a closed one, 𝒞^1D^ = 0, and this condition holds at the interface between the open and closed regions. For 𝒞^3D^, a nucleosome can interact with up to twelve nucleosomes in total, including two along the chromatin polymer and up to ten through 3D contacts. Since each of the ten 3D neighbors can be in an open (1) or closed (−1) state, 𝒞^3D^ takes an integer value from −10 to 10. The other parameters, *λ* and *ϕ*_*α*_ reflect the chromatin compaction and binding profile of the DNA-binding factor, respectively. Through the parameter *λ*, the chromatin 3D coordination {**R**_*n*_} affects the 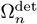, because *λ* is defined as the mean distance between three-dimensionally interacting nucleosomes around the *n*th nucleosome. Through the parameter *ϕ*_*α*_, DNA-binding factors affect the 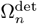, because *ϕ*_*α*_ is defined by their binding constants and binding sites distributions on the genome. Thus, the sign of opening determinant 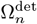, and hence that of 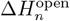, is determined by four epigenetic parameters, 𝒞^1D^, 𝒞^3D^, *λ*, and *ϕ*_*α*_.

### 2.3 Sequential opening condition

Next we investigated the sign of 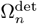, i.e., the direction of the state change of nucleosomes, on the (*ϕ*_*α*_, *λ*) plane. Sequential opening is defined as the path where the interface site *N*_int_ opens first, followed by *N*_int_+1, *N*_int_+2, … (figure 1 g), while no interior site opens prematurely. For any *N*_far_ *> N*_int_, premature opening of the *N*_far_th nucleosome is termed random opening. Using inequality (4), sequential opening persists when inequalities (12) hold simultaneously. To identify parameter regions satisfying inequalities (12), we plotted the curves 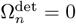 for *n* = *N*_int_ and *n* = *N*_far_ cases, for fixed (𝒞^1D^, 𝒞^3D^) (figure 2 e-h). The region with 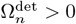 (to the right of the curves in our figures) denotes conditions where open state is stable. The overlap between the 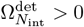 region and the 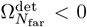 region highlights where sequential opening is favored (figure 2 e-h). Hereafter, we refer to this (*ϕ*_*α*_, *λ*) plot as a phase diagram.

**Figure 2.**
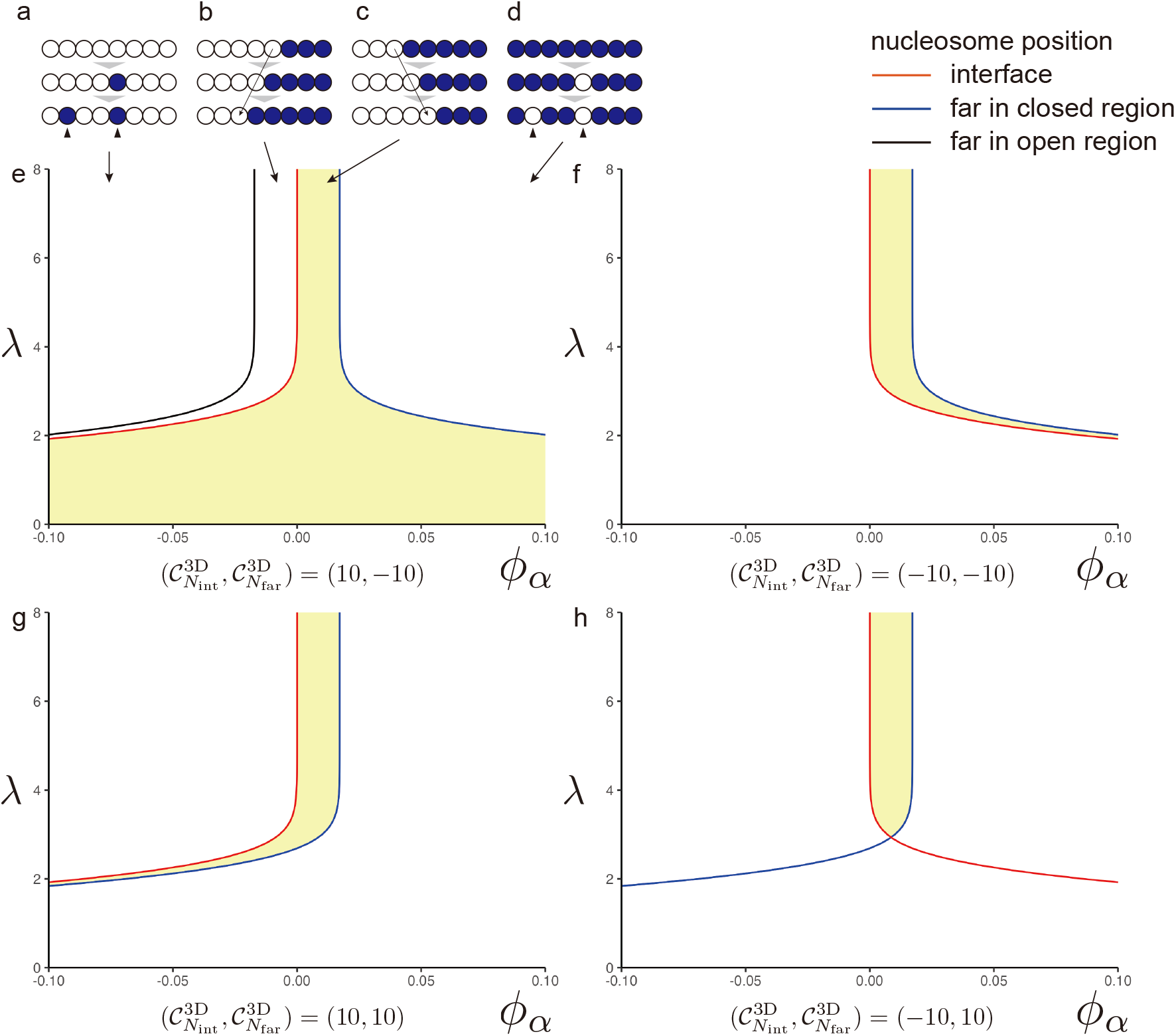
Opening order analyses. (a-d) opening or closing orders. (a) random closing, (b) sequential closing, (c) sequential opening that includes collinearity, highlighted in yellow in (e-h), and (d) random opening. (e-h) Phase diagrams of opening order parametrized by a binding profile of the DNA binding factor *ϕ*_*α*_ and the distance to three-dimensional neighbors,*λ*. (e) Interface is surrounded by open nucleosomes and the far is surrounded by closed nucleosomes. (f) Interface is surrounded by closed nucleosomes and the far is surrounded by closed nucleosomes. (g) Interface is surrounded by open nucleosomes and the far is surrounded by open nucleosomes. (h) Interface is surrounded by closed nucleosomes and the far is surrounded by open nucleosomes.

As an illustrative case, we considered the nucleosome coordination 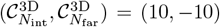 (figure 2 e), where the interface nucleosome is surrounded by 10 open neighbors and the *N*_far_th nucleosome is surrounded by 10 closed neighbors. The corresponding phase diagram (figure 2 e) reveals the following.

When *ϕ*_*α*_ is negative (closing bias), the interface opens only when *λ* is sufficiently small. This reflects a situation in which the interface nucleosome has nearby open neighbors in 3D interactions, forming a cluster that counteracts the closing bias. When *ϕ*_*α*_ is strongly positive (opening bias), *λ* must also be small for the *N*_far_th nucleosome to remain closed. In this case, the *N*_far_th nucleosome resides within a closed cluster, thereby resisting the opening bias. This is phenomenologically analogous to heterochromatin. Finally, with a moderately positive *ϕ*_*α*_ (mild opening bias), a sequential opening occurs for any *λ* under this coordination.

In general, the interface nucleosome requires the following conditions to be open. If surrounded by closed neighbors (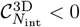, red curves in (figure 2 f, h)), opening occurs when *ϕ*_*α*_ is positive and *λ* is large, indicating that the opening bias of DNA-binding factors overcomes the inhibitory effects of closed neighbors (figure 2 f, h). If surrounded by open neighbors (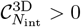, red curves in (figure 2 e, g)), opening occurs when *λ* is small, such that the strong local influence of open neighbors can dominate even under a closing bias (*ϕ*_*α*_ < 0) (figure 2 e, g).

For the *N*_far_th nucleosome, the following conditions are required to remain closed. If surrounded by closed neighbors (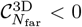, blue curves in (figure 2 e, f)), they remain closed when *ϕ*_*α*_ is small (mild opening bias), even with a small *λ* (dense nucleosome packing) (figure 2 e, f). If surrounded by open neighbors (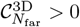, blue curves in (figure 2 g, h)), keeping a closed state requires a large *λ* (sparse nucleosome packing) and a sufficiently small *ϕ*_*α*_ (mild opening bias) (figure 2 g, h). Combining this condition for the *N*_far_th nucleosome to remain closed with another condition for the interface nucleosome to open, sequential opening arises when *ϕ*_*α*_ and *λ* are balanced appropriately across all types of nucleosome coordination (figure 2 e–h).

Conversely, our model characterizes the epigenetic conditions under which a non-collinear opening becomes energetically favorable. Outside the sequential opening region, the phase diagram (figure 2 e–h) predicts the conditions for (i) random opening, (ii) sequential closing, and (iii) random closing. Specifically, random opening occurs when 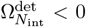 and 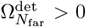. By symmetry, replacing the open and closed states, the conditions for sequential closing and random closing were revealed in the same way.

Overall, our theoretical framework provides a mechanistic explanation of how the order of chromatin opening is determined by the interplay between DNA-binding factor properties and nucleosome compaction.

## 3 Discussion

In this study, we theoretically investigated how the order of chromatin opening depends on the epigenetic state. Our model also revealed parameter regions in which sequential chromatin opening becomes energetically favorable.

Although many CREs and TFs have been reported to induce *Hox* gene expression at the appropriate position and timing, the mechanisms underlying the temporal expression order remain largely unknown [27]. Our theory suggests that the physical properties of chromatin polymers, particularly their interactions with DNA-binding factors, play a central role in determining the order of chromatin opening. The same physical mechanism could also account for sequential openings in other gene clusters [47–49], without invoking additional shared regulatory elements or specific TFs. Our model also predicts the existence of DNA-binding factors that randomly open nucleosomes (figure 2 e-h). Such factors may correspond to pioneer TFs that open nucleosomes, even in heterochromatin regions [65–69]. These factors exhibit high *ϕ*_*α*_ values and are located on the right side of the phase diagrams. Therefore, the binding profile *ϕ*_*α*_ can be interpreted as a pioneer tendency.

In this study, we focused on modeling the process by which TFs access DNA. To this end, we first introduced a chromatin polymer model in which each monomer takes multiple discrete states (see supplementary information for the details). For analytical tractability, we reduced the model to an Ising approximation with nucleosome state variable taking open and closed states. In physiological contexts, however, multiple epigenetic marks (e.g., DNA methylation, histone modifications) modulate chromatin, effectively altering nucleosome state stability and affinity of TFs [3–7, 9, 10]. Despite this simplification, our model provides a possible explanation for the sequential activation of the chromatin underlying *Hox* collinearity. An important direction for future research is to examine whether the present framework can be extended to capture complex and general epigenetic phenomena.

A limitation of our model is that several parameters introduced here are difficult to measure experimentally. Hi-C or microscopic imaging may enable the measurement of *λ*. Combining these results with ATAC-seq may enable the measurement of 𝒞^1D^ and 𝒞^3D^. For *ϕ*_*α*_, the binding profile of a DNA-binding factor, it is difficult to measure the affinity between DNA and the binding factor *in vivo*. The future development of measurement techniques and further theoretical derivations of easy-to-measure quantities are necessary.

Another limitation is that our theory focused only on energy-decreasing processes. However, research in nonequilibrium statistical mechanics has revealed that energy-increasing processes occur with low but finite probability. Our analysis ignored such low-probability events, under the assumption that they are kinetically suppressed and quickly relax back via energy-decreasing processes [70–75]. Nevertheless, there may be genomic regions where the reverse reaction does not occur immediately. Instead, a single nucleosome opens and is subsequently stabilized in the open state within a closed domain. This process is expected to occur in genomic regions where the conditions favor a net decrease in energy once the initial opening occurs. Regions enriched with TF binding sites with high *ϕ*_*α*_ binding profiles are predicted to be prone to this random openings. In our analysis of sequential opening, we assumed the presence of an open region and did not address how such an initial open region emerges. The emergence of this initial open region, including the opening events around the *Hox1* loci, is likely to require the random opening processes. In a physiological context, these regions are likely to function as enhancers. An important direction for future work is to examine whether the random opening processes predicted by our model can be observed at specific genomic loci. This can be addressed by combining ATAC-seq [76] and ChIP-based [77–80] experiments.

Overall, our framework provides a mechanistic basis for how nucleosome states and chromatin architecture set the order of chromatin opening and gene activation.

## 4 Materials and methods

### 4.1 Phase diagram plots

To plot phase diagrams, we solved the equation

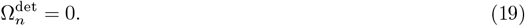

Using equations (10) and (19), we obtained the solution

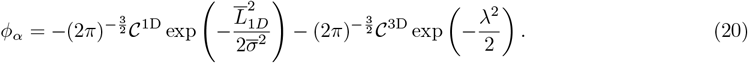

In the figure 2, we used the parameter ratio

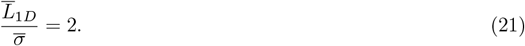

Additionally, 𝒞^1D^ = 0 for the interface and 𝒞^1D^ = − 2 for far from the interface. As illustrative cases, 𝒞^3D^ = 10 or 𝒞^3D^ = − 10 cases were plotted.

For details of the model definitions and approximations, see the supplementary information.

## 5 Acknowledgments

This research was supported by JSPS KAKENHI Grants (23K17000 to Y.A., JP19H05801 to Y.M., 25H01797 to T.S.) and by RIKEN Research Fund for Special Postdoctoral Researcher (Project Codes, 202401061040 and 202501094101).

## Author contributions

Y.A. and T.S. designed research; Y.A. and K.A. conducted theoretical analysis; T.S. and Y.M. offered important ideas; Y.A. wrote the original draft, and all authors reviewed and edited the manuscript.

## Data availability

All data and materials necessary to support the conclusions of this study are included in the article and the Supplementary Information.

