## Supplemental information for "A thermodynamic chromatin polymer model characterizes the epigenetic conditions for *Hox* collinearity"

**2 Relationships between energy, entropy, and probability 5**

### 1 Details for model definition and approximations

#### 1.1 Detailed model definition

We modeled chromatin as a polymer consisting of nucleosomes based on a previous model research [1], as monomers with Potts-like interactions between them, mediated by DNA-binding factors. The Hamiltonian reads

$$H = H_{\mathrm{GC}} + H_{\mathrm{VE}} + H_{\mathrm{F}} + H_{\mathrm{M}} + H_{\mathrm{P}}, \tag{1}$$

where

$$H_{\mathrm{GC}} = \frac{3k_BT}{2b^2}\sum_{n=1}^{N}\left(\mathbf{R}_n - \mathbf{R}_{n-1}\right)^2, \tag{2}$$

$$H_{\mathrm{VE}} = v\sum_{n\neq m}\mathcal{G}\left(\mathbf{R}_n - \mathbf{R}_m, \sigma\right) + w\sum_{n\neq m\neq k}\mathcal{G}\left(\mathbf{R}_n - \mathbf{R}_m, \sigma\right)\mathcal{G}\left(\mathbf{R}_n - \mathbf{R}_k, \sigma\right), \tag{3}$$

$$H_{\mathrm{F}} = -\mathbf{F}(\mathbf{R}_N - \mathbf{R}_0) \tag{4}$$

$$H_M = -\sum_{n}\sum_{\alpha,\beta}\sum_{\eta_\alpha}\pi(s_n,\alpha)\mathcal{G}\left(\mathbf{R}_n - \mathbf{R}_{\eta_\alpha}, \varsigma_\alpha\right)\mathbb{I}\left[n\in\mathcal{T}_\alpha\right], \tag{5}$$

$$\begin{aligned} H_P = &-\sum_{n,m}\sum_{\alpha,\beta}\sum_{\eta_\alpha,\zeta_\beta}\mathcal{K}(s_n,\alpha)\mathcal{K}(s_m,\beta)\mathcal{M}_{\alpha\beta} \\ &\mathcal{G}\left(\mathbf{R}_n - \mathbf{R}_{\eta_\alpha}, \varsigma_\alpha\right)\mathcal{G}\left(\mathbf{R}_m - \mathbf{R}_{\zeta_\beta}, \varsigma_\beta\right)\mathcal{G}\left(\mathbf{R}_{\eta_\alpha} - \mathbf{R}_{\zeta_\beta}, \frac{\varsigma_\alpha + \varsigma_\beta}{2}\right)\mathbb{I}\left[n\in\mathcal{T}_\alpha\right]\mathbb{I}\left[m\in\mathcal{T}_\alpha\right] \end{aligned} \tag{6}$$

$$\begin{aligned} \simeq &-\sum_{n,m}\sum_{\alpha,\beta}\sum_{\eta_\alpha,\zeta_\beta}\mathcal{K}(s_n,\alpha)\mathcal{K}(s_m,\beta)\mathcal{M}_{\alpha\beta} \\ &\mathcal{G}\left(\mathbf{R}_n - \mathbf{R}_{\eta_\alpha}, \varsigma_\alpha\right)\mathcal{G}\left(\mathbf{R}_m - \mathbf{R}_{\zeta_\beta}, \varsigma_\beta\right)\mathcal{G}\left(\mathbf{R}_n - \mathbf{R}_m, \sigma\right)\mathbb{I}\left[n\in\mathcal{T}_\alpha\right]\mathbb{I}\left[m\in\mathcal{T}_\alpha\right], \end{aligned} \tag{7}$$

respectively. For these models, we introduced an isotropic Gaussian distribution function

$$\mathcal{G}\left(\mathbf{x}, s\right) := \frac{1}{(2\pi)^{\frac{3}{2}}\ s^3}\exp\left(-\frac{|\mathbf{x}|^2}{2s^2}\right), \tag{8}$$

and parameters; $b$ denoting the Kuhn length (the size of the monomer); $v, w > 0$ denoting two- and three-body volume exclusion interaction coefficients; $\mathbf{F}$ denoting force applied on both end of the polymer; $\pi(s_n,\alpha)$ denoting an interaction coefficient between a DNA-binding factor $\alpha$ and a nucleosome in the state $s_n$; $\mathcal{K}(s_n,\alpha)$ denoting an interaction coefficient between a DNA-binding factor $\alpha$ and a nucleosome in the state $s_n$ which forms a bridge of two nucleosomes; $\mathcal{M}_{\alpha\beta}$ denoting an interaction coefficient between tow DNA-binding factors $\alpha$ and $\beta$; $\sigma$ denoting the radius of a monomer; $\varsigma_\alpha$ denoting the radius of a DNA-binding factor $\alpha$; respectively. For freely-jointed chains $b = 2\sigma$, and in general $\sigma \ll b$. The approximated $H_P$ in equation (7) approaches the same limit as $H_P$ in equation (6) while $\sigma$ and $\varsigma_\alpha$ approach 0. Therefore, we used equation (7) to define $H_P$ as follows. Here, $\mathcal{T}_\alpha$ denotes the target region of a DNA-binding factor $\alpha$ on the genome and

$$\mathbb{I}[\text{condition}] = \begin{cases} 1 & (\text{condition is True}) \\ 0 & (\text{condition is False}), \end{cases} \tag{9}$$

i.e. $\mathbb{I}[\text{condition}]$ is 1 when the condition in square brackets holds or is 0 when it does not. DNA-binding factor-related parameters were defined as

$$\pi(s_n,\alpha) = \sum_i \pi_{i,\alpha}\delta_{s_n,i}, \tag{10}$$

$$\mathcal{K}(s_n,\alpha) = \sum_i \mathcal{K}_{i,\alpha}\delta_{s_n,i}, \tag{11}$$

for Potts-like states $s_n \in [1, 2, \cdots, q]$. For Ising model analysis, $s_n \in [1(\text{open}), -1(\text{closed})]$. In this case,

$$\pi(s_n,\alpha) = p_\alpha s_n, \tag{12}$$

$$\mathcal{K}(s_n,\alpha) = \kappa_\alpha s_n, \tag{13}$$

and

$$\left(\pi_{\text{open},\alpha}, \pi_{\text{close},\alpha}\right) = \left(p_\alpha, -p_\alpha\right), \tag{14}$$

$$\left(\mathcal{K}_{\text{open},\alpha}, \mathcal{K}_{\text{close},\alpha}\right) = \left(\kappa_\alpha, -\kappa_\alpha\right). \tag{15}$$

For the Ising case analyses in the main text, we used these parameters. Using definitions (5), (7), (12) and (13) for the Ising case,

$$H_M = -\sum_n \sum_\alpha \sum_{\eta_\alpha} p_\alpha s_n \mathcal{G}\left(\mathbf{R}_n - \mathbf{R}_{\eta_\alpha}, \varsigma_\alpha\right) \mathbb{I}\left[n \in \mathcal{T}_\alpha\right], \tag{16}$$

$$H_P = -\sum_{n,m} \sum_{\alpha,\beta} \sum_{\eta_\alpha,\zeta_\alpha} \kappa_\alpha \kappa_\beta \mathcal{M}_{\alpha\beta} s_n s_m$$
$$\mathcal{G}\left(\mathbf{R}_n - \mathbf{R}_{\eta_\alpha}, \varsigma_\alpha\right) \mathcal{G}\left(\mathbf{R}_m - \mathbf{R}_{\zeta_\beta}, \varsigma_\beta\right) \mathcal{G}\left(\mathbf{R}_n - \mathbf{R}_m, \sigma\right) \mathbb{I}\left[n \in \mathcal{T}_\alpha\right] \mathbb{I}\left[m \in \mathcal{T}_\alpha\right]. \tag{17}$$

For the later calculations, we introduce a unit length $L_u$, and we derived dimensionless parameters:

$$\overline{b} := \frac{b}{L_u}, \tag{18}$$

$$\overline{\mathbf{R}} := \frac{\mathbf{R}}{L_u}, \tag{19}$$

$$\overline{\sigma} := \frac{\sigma}{L_u}, \tag{20}$$

$$\overline{\varsigma_\alpha} := \frac{\varsigma_\alpha}{L_u}, \tag{21}$$

$$\overline{p}_\alpha := \frac{p_\alpha}{k_B T L_u^3}, \tag{22}$$

$$\overline{\kappa}_\alpha := \frac{\kappa_\alpha}{k_B T L_u^3}, \tag{23}$$

$$\overline{\mathcal{M}}_{\alpha\beta} := \frac{k_B T \mathcal{M}_{\alpha\beta}}{L_u^3}. \tag{24}$$

An enough large unit length $L_u$ gives the thermodynamic limit discussed previously [1]. Hereafter, we use the Kuhn length $b$ as the unit length, i.e. $L_u = b$.

Additionally, we introduce a dimensionless length between $n$th and $m$th nucleosomes,

$$\overline{L}_{nm} := \left|\overline{\mathbf{R}}_n - \overline{\mathbf{R}}_m\right|. \tag{25}$$

Only when $\alpha = \beta$, we introduced an abbreviation,

$$\overline{\mathcal{M}}_\alpha := \overline{\mathcal{M}}_{\alpha\alpha}. \tag{26}$$

#### 1.2 Detailed approximation of the opening determinant

As discussed in the main text, $H_{\mathrm{GC}}$, $H_{\mathrm{VE}}$ and $H_{\mathrm{F}}$ are state-independent, because of each definition. And therefore the difference is

$$\Delta H_n^{\mathrm{open}} := H\left(\{\mathbf{R}_m\}, \{s_m\}\right)\big|_{s_n=1} - H\left(\{\mathbf{R}_m\}, \{s_m\}\right)\big|_{s_n=-1}. \tag{27}$$

Then,

$$\frac{\Delta H_n^{\mathrm{open}}}{k_B T} = -(1-(-1)) \sum_m \Bigg[\overline{p}_\alpha \mathbb{I}\left[n \in \mathcal{T}_\alpha\right] \left(\sum_{\eta_\alpha} \mathcal{G}\left(\left|\overline{\mathbf{R}}_n - \overline{\mathbf{R}}_{\eta_\alpha}\right|, \overline{\varsigma}_\alpha\right)\right)$$
$$+ \overline{\kappa}_\alpha^2 \overline{\mathcal{M}}_\alpha s_m \mathbb{I}\left[n \in \mathcal{T}_\alpha\right] \mathbb{I}\left[m \in \mathcal{T}_\alpha\right] \mathcal{G}\left(\overline{L}_{nm}, \overline{\sigma}\right) \left(\sum_{\eta_\alpha,\zeta_\alpha} \mathcal{G}\left(\left|\overline{\mathbf{R}}_n - \overline{\mathbf{R}}_{\eta_\alpha}\right|, \overline{\varsigma}_\alpha\right) \mathcal{G}\left(\left|\overline{\mathbf{R}}_m - \overline{\mathbf{R}}_{\zeta_\beta}\right|, \overline{\varsigma}_\alpha\right)\right)\Bigg]$$
$$= -2\left[\overline{p}_\alpha \mathbb{I}_{n,\alpha} \mathbb{P}_{n,\alpha} + \sum_m \overline{\kappa}_\alpha^2 \overline{\mathcal{M}}_\alpha s_m \mathbb{I}_{n,\alpha} \mathbb{I}_{m,\alpha} \mathcal{G}\left(\overline{L}_{nm}, \overline{\sigma}\right) \mathbb{P}_{n,\alpha} \mathbb{P}_{m,\alpha}\right] \tag{28}$$
$$= -2 \mathbb{I}_{n,\alpha} \mathbb{P}_{n,\alpha} \overline{\kappa}_\alpha^2 \overline{\mathcal{M}}_\alpha \Omega_n^{\mathrm{det}}, \tag{29}$$

where the opening determinant was defined as,

$$\Omega_n^{\mathrm{det}} := \frac{\overline{p}_\alpha}{\overline{\kappa}_\alpha^2 \overline{\mathcal{M}}_\alpha} + \sum_m s_m \mathcal{G}\left(\overline{L}_{nm}, \overline{\sigma}\right) \mathbb{I}_{m,\alpha} \mathbb{P}_{m,\alpha}. \tag{30}$$

The inter-nucleosome effects can be divided into 1D and 3D effects as,

$$\Omega_n^{\mathrm{det}} = \frac{\overline{p}_\alpha}{\overline{\kappa}_\alpha^2 \overline{\mathcal{M}}_\alpha} + \sum_{m \in [n \pm 1]} s_m \mathcal{G}\left(\overline{L}_{nm}, \overline{\sigma}\right) \mathbb{I}_{m,\alpha} \mathbb{P}_{m,\alpha} + \sum_{m \notin [n \pm 1]} s_m \mathcal{G}\left(\overline{L}_{nm}, \overline{\sigma}\right) \mathbb{I}_{m,\alpha} \mathbb{P}_{m,\alpha}. \tag{31}$$

Because the $m$th nucleosomes here are sufficiently close to the $n$th one to contact it, we approximated that the probability that the $n$-nucleosome contacts a TF $\alpha$ is the same as the probability that the geometrically neighboring $m$th nucleosome contacts the TF $\alpha$, that is, $\mathbb{P}_{m,\alpha} = \mathbb{P}_{n,\alpha}$. As another approximation, we replaced the target identity of TF $\alpha$ on the $m$th nucleosome, $\mathbb{I}_{m,\alpha}$, with its genomic density $\mathbb{D}_\alpha$. Then,

$$\Omega_n^{\mathrm{det}} \simeq \mathbb{D}_\alpha \mathbb{P}_{n,\alpha} \left[ \frac{\overline{p}_\alpha}{\mathbb{D}_\alpha \mathbb{P}_{n,\alpha} \overline{\kappa}_\alpha^2 \overline{\mathcal{M}}_\alpha} + \sum_{m \in [n\pm1]} s_m \mathcal{G}\left(\overline{L}_{nm}, \overline{\sigma}\right) + \sum_{m \notin [n\pm1]} s_m \mathcal{G}\left(\overline{L}_{nm}, \overline{\sigma}\right) \right]. \tag{32}$$

Taking the mean, the distance from the $m$th nucleosome to the $n$th nucleosome can be approximated to be the same. Then,

$$\Omega_n^{\mathrm{det}} \simeq \mathbb{D}_\alpha \mathbb{P}_{n,\alpha} \left[ \frac{\overline{p}_\alpha}{\mathbb{D}_\alpha \mathbb{P}_{n,\alpha} \overline{\kappa}_\alpha^2 \overline{\mathcal{M}}_\alpha} + \mathcal{G}\left(\overline{L}_{1\mathrm{D}}, \overline{\sigma}\right) \sum_{m \in [n\pm1]} s_m + \mathcal{G}\left(\overline{L}_{3\mathrm{D}}, \overline{\sigma}\right) \sum_{m \notin [n\pm1]} s_m \right] \tag{33}$$

$$= \mathbb{D}_\alpha \mathbb{P}_{n,\alpha} \left[ \frac{\overline{p}_\alpha}{\mathbb{D}_\alpha \mathbb{P}_{n,\alpha} \overline{\kappa}_\alpha^2 \overline{\mathcal{M}}_\alpha} + \mathcal{C}^{1\mathrm{D}} \mathcal{G}\left(\overline{L}_{1\mathrm{D}}, \overline{\sigma}\right) + \mathcal{C}^{3\mathrm{D}} \mathcal{G}\left(\overline{L}_{3\mathrm{D}}, \overline{\sigma}\right) \right]. \tag{34}$$

where

$$\mathcal{C}^{1\mathrm{D}} = \sum_{m \in [n\pm1]} s_m \tag{35}$$

$$\mathcal{C}^{3\mathrm{D}} = \sum_{m \notin [n\pm1]} s_m. \tag{36}$$

By defining two key parameters, the distance in 3D contact and binding profile of the DNA-binding factor, as

$$\lambda := \frac{\overline{L}_{3D}}{\overline{\sigma}}, \tag{37}$$

$$\phi_\alpha := \frac{\overline{\sigma}^3 \overline{p}_\alpha}{\mathbb{D}_\alpha \mathbb{P}_{n,\alpha} \overline{\kappa}_\alpha^2 \overline{\mathcal{M}}_\alpha}, \tag{38}$$

the opening determinant was derived as

$$\Omega_n^{\mathrm{det}} \simeq \frac{\mathbb{D}_\alpha \mathbb{P}_{n,\alpha}}{\overline{\sigma}^3} \left[ \phi_\alpha + \overline{\sigma}^3 \mathcal{C}^{1\mathrm{D}} \mathcal{G}\left(\overline{L}_{1\mathrm{D}}, \overline{\sigma}\right) + \overline{\sigma}^3 \mathcal{C}^{3\mathrm{D}} \mathcal{G}\left(\overline{L}_{3\mathrm{D}}, \overline{\sigma}\right) \right] \tag{39}$$

$$= \frac{\mathbb{D}_\alpha \mathbb{P}_{n,\alpha}}{(2\pi)^{\frac{3}{2}} \overline{\sigma}^3} \left[ (2\pi)^{\frac{3}{2}} \phi_\alpha + \mathcal{C}^{1\mathrm{D}} \exp\left( -\frac{\overline{L}_{1\mathrm{D}}^2}{2\overline{\sigma}^2} \right) + \mathcal{C}^{3\mathrm{D}} \exp\left( -\frac{\lambda^2}{2} \right) \right]. \tag{40}$$

Here, we used the definition of Gaussian distribution function in equation (8). Then, with the equation (29), i.e., $\Delta H_n^{\mathrm{open}}/k_B T = -2\mathbb{I}_{n,\alpha}\mathbb{P}_{n,\alpha}\overline{\kappa}_\alpha^2 \overline{\mathcal{M}}_\alpha \Omega_n^{\mathrm{det}}$, parameter $(\mathcal{C}^{1\mathrm{D}}, \mathcal{C}^{3\mathrm{D}}, \lambda, \phi_\alpha)$ region which gives $\Omega_n^{\mathrm{det}} > 0$ decreases the total energy by opening the $n$th nucleosome. We further analyzed the opening determinant $\Omega_n^{\mathrm{det}}$ in the main text.

#### 1.3 $\alpha \neq \beta$ and $\overline{\kappa}_\alpha \overline{\kappa}_\beta \overline{\mathcal{M}}_{\alpha\beta} < 0$ cases

When not necessarily $\alpha = \beta$, equation (27) was expanded into,

$$\frac{\Delta H_n^{\mathrm{open}}}{k_B T} = -2 \left[ \overline{p}_\alpha \mathbb{I}_{n,\alpha} \mathbb{P}_{n,\alpha} + \sum_m \overline{\kappa}_\alpha \overline{\kappa}_\beta \overline{\mathcal{M}}_{\alpha\beta} s_m \mathbb{I}_{n,\alpha} \mathbb{I}_{m,\beta} \mathcal{G}\left(\overline{L}_{nm}, \overline{\sigma}\right) \mathbb{P}_{n,\alpha} \mathbb{P}_{m,\beta} \right], \tag{41}$$

instead of equation (28). Then, using the same definitions of approximated parameters $\mathcal{C}^{1\mathrm{D}}, \mathcal{C}^{3\mathrm{D}}$, and $\mathbb{D}_\alpha$,

$$\frac{\Delta H_n^{\mathrm{open}}}{k_B T} \simeq \frac{\overline{\kappa}_\alpha \overline{\kappa}_\beta \overline{\mathcal{M}}_{\alpha\beta} C^{(-)}}{\overline{\sigma}^3} \left[ \phi_{\alpha\beta} + \overline{\sigma}^3 \mathcal{C}^{1\mathrm{D}} \mathcal{G}\left(\overline{L}_{1\mathrm{D}}, \overline{\sigma}\right) + \overline{\sigma}^3 \mathcal{C}^{3\mathrm{D}} \mathcal{G}\left(\overline{L}_{3\mathrm{D}}, \overline{\sigma}\right) \right], \tag{42}$$

where an always-negative coefficient $C^{(-)}$ was defined as

$$C^{(-)} := -2\mathbb{D}_\alpha \mathbb{D}_\beta \mathbb{P}_\alpha \mathbb{P}_\beta \tag{43}$$

and a binding profile parameter of two TFs indexed by $\alpha$ and $\beta$ as

$$\phi_{\alpha\beta} := \frac{\overline{\sigma}^3 \, \overline{p}_\alpha}{\mathbb{D}_\alpha \mathbb{D}_\beta \mathbb{P}_\beta \overline{\kappa}_\alpha \overline{\kappa}_\beta \overline{\mathcal{M}}_{\alpha\beta}}. \tag{44}$$

This $\phi_{\alpha\beta}$ definition is a generalization of $\phi_\alpha$ in the equation (38), for a case where two different DNA-binding factors bridge two DNA regions. When $\overline{\kappa}_\alpha\overline{\kappa}_\beta\overline{\mathcal{M}}_{\alpha\beta} > 0$, using the equation (42) and (8), an inequality

$$\phi_{\alpha\beta} + \overline{\sigma}^3\mathcal{C}^{1\mathrm{D}}\mathcal{G}\left(\overline{L}_{1\mathrm{D}}, \overline{\sigma}\right) + \overline{\sigma}^3\mathcal{C}^{3\mathrm{D}}\mathcal{G}\left(\overline{L}_{3\mathrm{D}}, \overline{\sigma}\right) > 0 \tag{45}$$

gives the condition of energy decrease by $n$th nucleosome opening. It is the same condition as $\Omega_n^{\mathrm{det}} > 0$ with the $\Omega_n^{\mathrm{det}}$ definition by equation (40). When $\overline{\kappa}_\alpha\overline{\kappa}_\beta\overline{\mathcal{M}}_{\alpha\beta} < 0$, an inequality

$$\phi_{\alpha\beta} + \overline{\sigma}^3\mathcal{C}^{1\mathrm{D}}\mathcal{G}\left(\overline{L}_{1\mathrm{D}}, \overline{\sigma}\right) + \overline{\sigma}^3\mathcal{C}^{3\mathrm{D}}\mathcal{G}\left(\overline{L}_{3\mathrm{D}}, \overline{\sigma}\right) < 0 \tag{46}$$

gives the condition of energy decrease by $n$th nucleosome opening. Then $\Delta H_{N_{\mathrm{int}}}^{open} < 0$ and $\Delta H_{N_{\mathrm{far}}}^{open} > 0$ region is the highlighted area in the Figure S1.

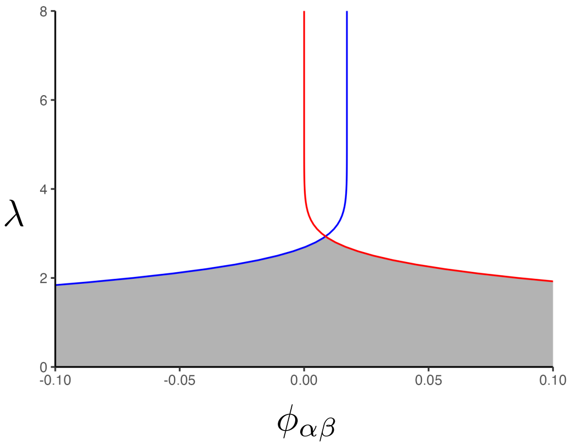

Figure S1. A phase diagram in $\overline{\kappa}_\alpha\overline{\kappa}_\beta\overline{\mathcal{M}}_{\alpha\beta} < 0$ case. Only when the interface is surrounded by closed nucleosomes ($\mathcal{C}_{N_{\mathrm{int}}}^{3\mathrm{D}} < 0$, here $\mathcal{C}_{N_{\mathrm{int}}}^{3\mathrm{D}} = -10$) and the far is surrounded by open nucleosomes ($\mathcal{C}_{N_{\mathrm{far}}}^{3\mathrm{D}} > 0$, here $\mathcal{C}_{N_{\mathrm{far}}}^{3\mathrm{D}} = 10$), there is the parameter region in which the sequential opening is energetically favored in $\overline{\kappa}_\alpha\overline{\kappa}_\beta\overline{\mathcal{M}}_{\alpha\beta} < 0$ condition.

### 2 Relationships between energy, entropy, and probability

In the main text, we analyzed the energy changes. Based on the Boltzmann distribution in the equilibrium state,

$$p(\{\mathbf{R}_n\}, \{s_n\}) = Z^{-1}\exp\left(\frac{-H(\{\mathbf{R}_n\}, \{s_n\})}{k_B T}\right), \tag{47}$$

$$Z = \sum_{\{\mathbf{R}_n\},\{s_n\}} \exp\left(\frac{-H(\{\mathbf{R}_n\}, \{s_n\})}{k_B T}\right) \tag{48}$$

where $k_B, T$, $H(\{\mathbf{R}_n\}, \{s_n\})$, and $Z$ indicate the Boltzmann constant, temperature, the Hamiltonian, which represents the total energy of the chromatin, and the partition function, respectively. The partition function $Z$ is a normalization coefficient that ensures that the sum of the probabilities is equal to one. Moreover, it is known that the Helmholtz free energy $\mathcal{F}$ is derived from $Z$ as,

$$\mathcal{F} = k_B T \ln Z. \tag{49}$$

Because it is usually difficult to solve the integrations within $Z$, it is also difficult in general to analytically solve the Helmholtz free energy. Therefore, a pseudo free energy $f$ taking arguments $x$ is defined as a function such that

$$Z = \int_{\forall x} dx \exp\left(-\frac{f(x)}{k_B T}\right), \tag{50}$$

and it is analyzed to investigate the effective energy of the system. Previous studies [1, 2] have indicated that the balance between energetic and entropic contributions varies across conditions. In this study, we focused on dense configurations of the chromatin polymer, in which the energetic term was reported to dominate [1]. Therefore, we restrict our analysis to the energetic contribution in the main text.
